# SeqTagger, a rapid and accurate tool to demultiplex direct RNA nanopore sequencing datasets

**DOI:** 10.1101/2024.10.29.620808

**Authors:** Leszek P Pryszcz, Gregor Diensthuber, Laia Llovera, Rebeca Medina, Anna Delgado-Tejedor, Luca Cozzuto, Julia Ponomarenko, Eva Maria Novoa

## Abstract

Nanopore direct RNA sequencing (DRS) enables direct measurement of RNA molecules, including their native RNA modifications, without prior conversion to cDNA. However, commercial methods for molecular barcoding of multiple DRS samples are lacking, and community-driven efforts, such as DeePlexiCon, are not compatible with newer RNA chemistry flowcells and the latest-generation GPU cards. To overcome these limitations, we introduce SeqTagger, a rapid and robust method that can demultiplex direct RNA sequencing datasets with 99% precision and 95% recall. We demonstrate the applicability of SeqTagger in both RNA002/R9.4 and RNA004/RNA chemistries and show its robust performance both for long and short RNA libraries, including custom libraries that do not contain standard poly-(A) tails, such as Nano-tRNAseq libraries. Finally, we demonstrate that increasing the multiplexing up to 96 barcodes yields highly accurate demultiplexing models. SeqTagger can be executed in a standalone manner or through the MasterOfPores NextFlow workflow. The availability of an efficient and simple multiplexing strategy improves the cost-effectiveness of this technology and facilitates the analysis of low-input biological samples.

## INTRODUCTION

Nanopore sequencing technologies have revolutionised our ability to study the transcriptome, by enabling direct sequencing of the native RNA molecules, offering insights into gene expression profiles while retaining RNA modification and polyA tail length information (Garalde et al. 2018). Direct RNA sequencing offers several advantages over traditional sequencing methods such as obviating the need for PCR amplification and cDNA synthesis steps, thus preserving the native state of RNA molecules and providing a less biassed and comprehensive view of the transcriptome (Workman et al. 2019).

However, key challenges persist, such as the lack of commercial barcoding kits and demultiplexing algorithms specific to direct RNA sequencing (DRS) data, hampering the broader use of this technology. To overcome these limitations, community-driven efforts have been made to enable demultiplexing of DRS data, such as PorePlex or DeePlexiCon (Smith et al. 2020). DeePlexiCon relies on the conversion of signal information to images, which are then classified using 2D convolutional neural networks (2D CNNs) into their corresponding barcodes. This is a computationally expensive task in the bioinformatic processing of direct RNA-sequencing data (Cozzuto et al. 2023). In addition, the user has to decide whether high demultiplexing accuracy or recovery takes precedence, with DeePlexiCon being able to reach 99% accuracy sacrificing around 40% of total reads or 92% accuracy losing 7% of reads (Smith et al. 2020).

Here we reasoned that developing a novel demultiplexing software that relies on basecalling the reverse transcription adapter (RTA), used during direct RNA-sequencing library preparation, should yield superior performance compared to existing tools. We rigorously benchmarked this novel tool, which we named *SeqTagger*, against the *de-facto* standard *DeePlexiCon*, to determine both improvements in model performance (recall and precision) and system performance (computation time and CPU usage). The applicability of *SeqTagger* was tested across sequencing devices (MinION and PromethION), sequencing chemistries (SQK-RNA002 and SQK-RNA004), RNA biotypes (poly-(A)^+^ and tRNA), and for extended sets of barcodes (up to 96 barcodes).

## RESULTS

### Demultiplexing DRS libraries through direct basecalling of the DNA barcode

To date, efforts for demultiplexing DRS datasets have relied on the transformation of signal intensities to images, followed by classification of resulting 2D images, as performed by the software DeePlexiCon (Smith et al. 2020). While this approach leads to robust and accurate demultiplexing statistics, it is computationally expensive, limited to 4 barcodes, and does not support newer RNA004 kit chemistries.

We hypothesised that training a DNA basecaller that could directly basecall a DNA barcode sequence present in the reverse transcription adapter (RTA), which is ligated to the native RNA molecules during library preparation, should lead to improved performance of the demultiplexing algorithm. In particular, the use of a DNA basecaller would allow skipping the most computationally expensive step of demultiplexing, i.e. the signal transformation step (Smith et al. 2020), and would in principle be flexible towards classifying an unlimited number of barcodes with minimal losses in demultiplexing accuracy, thus allowing to increase the number of samples that are loaded in a single flowcell.

We should note that the need for training a new DNA basecalling model – rather than using existing pre-trained DNA basecalling models – arises from the fact that the DNA barcode is sequenced using the ‘RNA’ chemistry, which exhibits several key differences to the DNA chemistry, preventing the use existing DNA basecalling models : i) RNA is sequenced from its 3’ end (in 3’>5’ direction), while DNA is sequenced from its 5’ end (in 5’>3’ direction); ii) the RNA helicase is slower than the DNA helicase (70 bps for RNA002 and 130 bps for RNA004, compared to 400-450 bps for DNA); and iii) RNA molecules are sampled with a different frequency than DNA molecules (3-4kHz for RNA, compared to 4-5kHz for DNA, depending on the flowcell). Thus, the same DNA barcode will generate different signals when sequenced with DNA and RNA kits, making existing DNA basecalling models unusable for basecalling of DNA barcodes in DRS datasets.

To demultiplex reads, SeqTagger performs several consecutive steps on each read: (i) signal segmentation (trimming of barcode signal) and normalisation, (ii) decoding of the signal into the sequence space (basecalling) followed by mapping to the reference, and (iii) filtering out potential misassigned barcodes using per-read median base quality information (**Fig. 1**). To train the basecalling model, we chose the latest CTC-CRF model architecture that is achieving over 99% basecalling accuracy for DNA basecalling (https://nanoporetech.com/platform/accuracy/). Training was performed using the *Bonito* software (https://github.com/nanoporetech/bonito/) (see *Methods*), and trained models (listed in **Supplemental Table S1**) were independently validated on PromethION direct RNA sequencing datasets, both for RNA002 and RNA004 chemistries (see **Supplemental Table S2** for full list of DRS datasets used).

**Figure 1.**
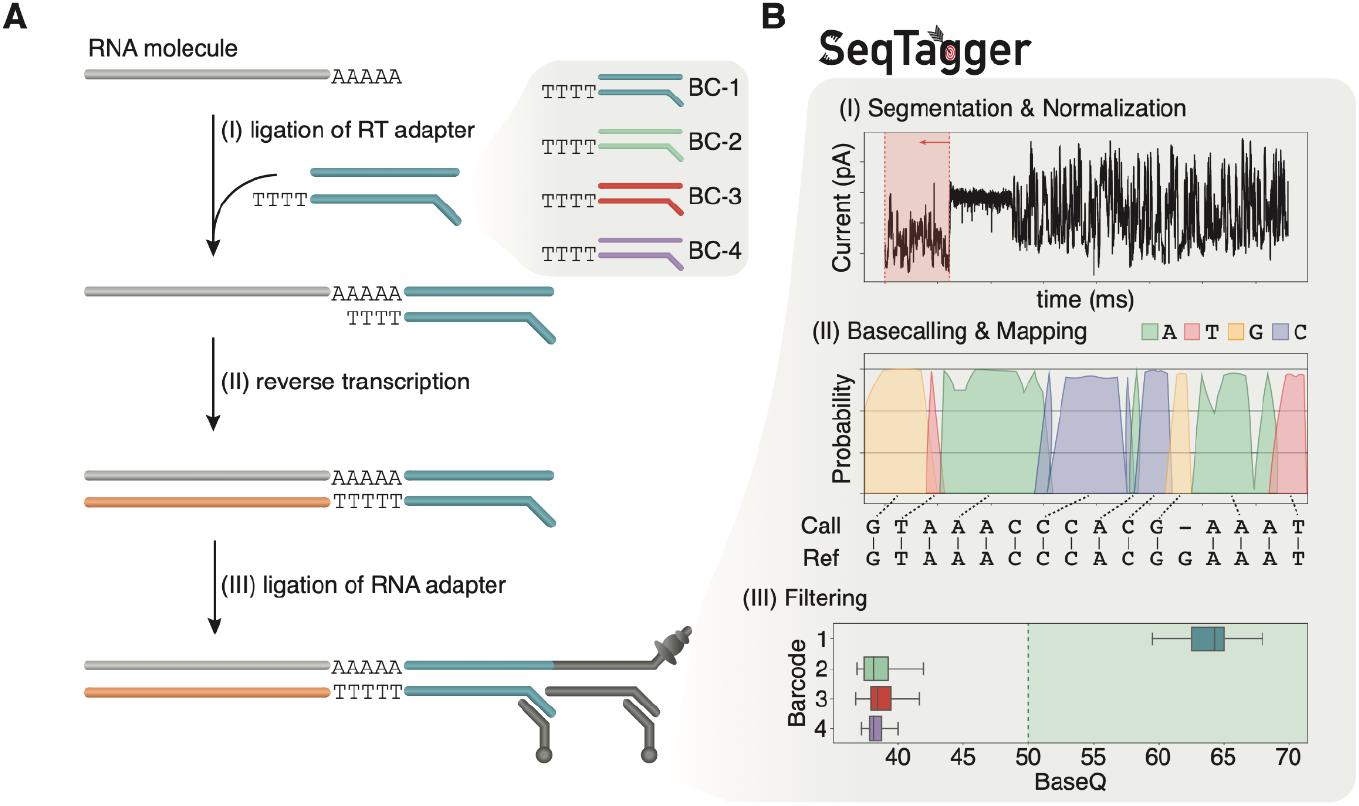
Schematic overview of the direct RNA sequencing (DRS) demultiplexing workflow. **(A)** Overview of the barcoded direct RNA sequencing workflow in which the standard reverse transcription adapter (RTA) is replaced with a barcode-containing adapter. Following adapter ligation and reverse transcription the RNA ligation adapter (RLA), containing the helicase enzyme, is ligated, making the library sequencing-ready. **(B)** Overview of the demultiplexing workflow performed by SeqTagger. The algorithm first segments the raw current intensity signal by identifying the poly-(A)-tail signal to extract the barcode-containing RT adapter. Following signal normalisation, the DNA sequence is basecalled and aligned to a set of reference barcodes. Finally, a filtering step is applied based on the median base quality (BaseQ) to remove misassigned barcode sequences.

### Systematic benchmarking of DRS demultiplexing software

We first trained SeqTagger on DRS data sequenced with R9.4/RNA002 chemistry, using 4 different barcodes that were embedded within the RT adapter (**Fig. 1A**). Notably, barcode sequences were kept identical to those used to train the DeePlexiCon model, to allow for direct comparison between SeqTagger and DeePlexiCon (see **SupplementalTable S1** and *Methods*).

To test the performance of the trained models, we generated three datasets of 100,000 reads each from an independently sequenced sample, in which each barcode had been ligated to a different *in vitro* transcribed (IVT) RNA (**Supplemental Table S2**). In this design, the ground truth is known thus allowing us to determine classification metrics of the trained model (see **Fig. 2A** and **Supplemental Table S3**). We should note that this independently sequenced run was not used for training/validation or of either demultiplexing tool. To facilitate the monitoring of resources, and ensure that the computational resources used by each of the software were directly comparable, the data was analysed using MasterOfPores (Cozzuto et al. 2023; Di Tommaso et al. 2017), a nextflow workflow for the analysis of DRS data, with the option ‘SeqTagger’ (model b04_RNA002), ‘DeePlexiCon’ (model resnet20-final.h5) or ‘no demultiplexing’ (ground-truth) (see *Methods*).

**Figure 2.**
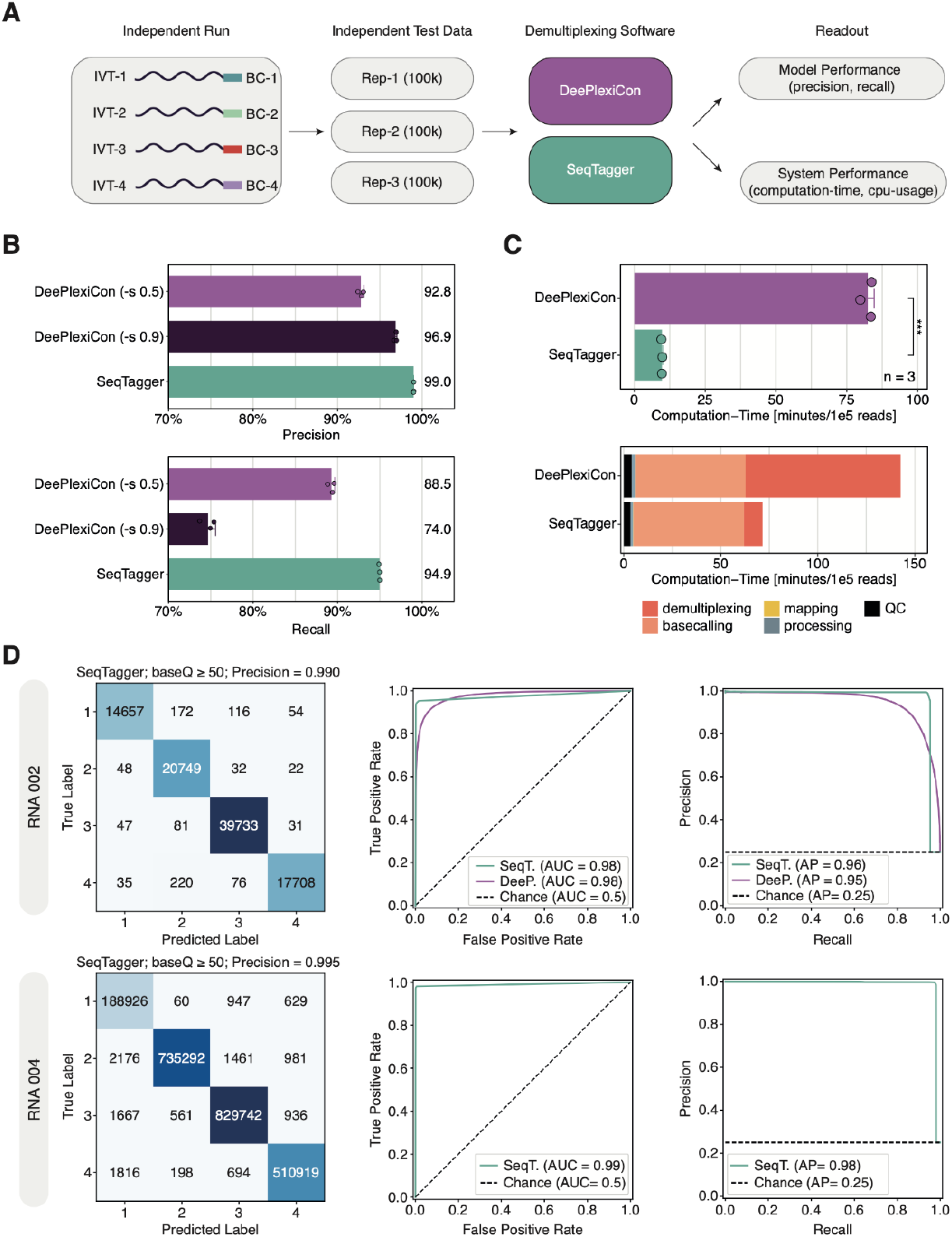
Comprehensive benchmarking of SeqTagger performance. **(A)** Schematic overview of the workflow used for comparative analysis of DRS demultiplexing software: DeePlexicon (purple) and SeqTagger (green). **(B)** Barplots depicting the demultiplexing precision and recall achieved with SeqTagger default settings (baseQ > 50), DeePlexiCon high recovery settings (-s 0.5), and DeePlexiCon high accuracy settings (-s 0.9), on the same 3 datasets described in panel A. Bars represent the mean (also indicated by the numeric value to the right of each bar) with error bars showing +/-1 standard deviation. Dots represent individual replicates. **(C)** Top: Barplot depicting the computation time of SeqTagger and DeePlexiCon, on the benchmarking datasets. Bars represent the mean value with error bars indicating +/-1 standard deviation. Dots represent individual replicates. Statistical significance was determined using a two-sided t-test (ns: p > 0.05, *: p <= 0.05, **: p <= 0.01, ***: p <= 0.001). Bottom: Barplot representing the absolute contribution of individual preprocessing steps to the total computation time (Rep-1). **(D)** Confusion matrices (left), ROC curves (middle), and Precision-Recall curves (right) on independent test data generated with RNA002 and RNA004 kit chemistries. Data was analysed with SeqTagger model b04_RNA002 (upper panels) and b04_RNA004 (bottom panels), respectively. Abbreviations: AUC (Area Under the Curve); AP (Average Precision).

We then compared SeqTagger results (using default settings, i.e. baseQ-cutoff ≥ 50) to those obtained with DeePlexiCon, either using settings for high recovery (-s 0.5) or high accuracy (-s 0.9). Samples were processed using GPU computing nodes running CUDA10, as CUDA11 is not supported by DeePlexiCon, and thus would not be directly comparable to SeqTagger. Of note, SeqTagger can be executed both in GPU computing nodes running CUDA10 and CUDA11. Our analysis revealed that SeqTagger reached 99% precision on the independent test datasets, whereas DeePlexiCon achieved 93% and 97% precision with the high recovery and high accuracy settings, respectively (**Fig. 2B**, top panel, *see also* **Supplemental Table S3**). In addition, SeqTagger achieved a recall of 95%, whereas DeePlexiCon’s high recovery mode reached 89%, followed by the high accuracy mode with 75% recall (**Fig. 2B**, bottom panel, *see also* **Supplemental Table S3**).

Next, we examined the computing time and resources required for demultiplexing. Our analysis demonstrated that SeqTagger was ∼9x faster than DeePlexiCon (**Fig. 2C** top panel, **Supplemental Fig. S1A**, and **Supplemental Table S4**). Notably, this increase in speed has important implications for the pre-processing of DRS data, as the demultiplexing step is typically a major computational bottleneck, taking up ≥ 50% of the overall computation time (**Fig. 2C** bottom panel, **Supplemental Fig. S1B**, and **Supplemental Table S4**). Similar results were observed when performing the same analysis on a more complex dataset of mouse poly(A)-selected material, where DeePlexiCon was found to take ∼40% of the overall computation time while SeqTagger reduced this to 8.5% of the total time required (**Supplemental Fig. S1C**).

Finally, we tested SeqTagger’s classification performance on *in vivo* data, by using two runs of poly(A)-tailed, total RNA from *E. coli* (Delgado-Tejedor et al. 2023). Both runs contained two (out of the four possible) barcodes used in our b04 models, thus allowing us to determine the false positive rate per barcode. This showed that between 0.1-0.2% of total demultiplexed reads were incorrectly assigned per barcode using SeqTagger, resulting in an overall precision of ≥ 99% (**Supplemental Fig. S1D**, *see also* **Supplemental Table S5)**. By contrast, DeePlexiCon had higher amounts of incorrectly predicted barcodes, reaching 3-6% per barcode in high-recovery mode and 1-3% in high-accuracy mode (**Supplemental Fig. S1D)**. In addition, SeqTagger recovered higher proportions of demultiplexed reads (90-93% recall) than DeePlexiCon under high-recovery (81-88% recall) or high-accuracy modes (65-73% recall), supporting our observations made on *in vitro* datasets (*see* **Supplemental Table S5**).

Taken together, these results demonstrate that demultiplexing direct RNA sequencing datasets by basecalling the DNA part of the RT adapter yields superior performance both in terms of precision and recall, while providing a significant increase in speed over existing tools.

### SeqTagger is compatible with both RNA002 and RNA004 chemistries

We then examined whether SeqTagger would be able to demultiplex DRS datasets sequenced with the recently upgraded chemistry that ONT has released for sequencing DRS datasets, which uses SQK-RNA004 kits (replacing SQK-RNA002 kits) and ‘RNA’ flowcells (replacing R9.4 flowcells). Notably, demultiplexing options are currently unavailable for this newer chemistry, neither commercially nor from the scientific community, thus making the transition to the newer chemistry highly problematic.

Here, we prepared DRS libraries using the recently released SQK-RNA004 library preparation kit on the same 4 barcodes (**Table S2**) ligated to IVT products as previously described, and trained a new demultiplexing model using *Bonito* (b04_RNA004). The performance of the model was assessed using reads from an independent test dataset (flowcell not used for training/validation of the model), as previously done for the SQK-RNA002/R9.4 chemistry (**Fig. 2D** top panels, **Supplemental Fig. S1E**, and **Supplemental Table S3**). Our results showed that SeqTagger was able to reach an overall precision above 99% with a recall of 97% on the new RNA004 chemistry (**Supplemental Fig. 2D** bottom panels), thus showing improved performance compared to the old R9.4/RNA002 chemistry.

Altogether, our results demonstrate that SeqTagger is a robust DRS demultiplexing algorithm, reaching ≥ 99% precision on both RNA002 and RNA004 chemistries while recovering 95% and 97% of all reads, respectively. Notably, the increased demultiplexing speed that direct basecalling of the DNA barcode offers (**Fig. 2C**), makes SeqTagger well-suited for larger-sized datasets that are expected to be generated with RNA004 kits (due to increased helicase speed and likely higher flowcell longevity), thus removing a potential computational bottleneck in DRS data analyses.

### SeqTagger can be extended to different RNA biotypes

Commercial DRS library preparation kits were initially designed to sequence poly-(A)-tailed RNA (mRNA, lncRNA, etc.) or *in-vitro* polyadenylated RNA molecules. For this reason, many DRS-related software, such as DeePlexiCon, were trained on polyadenylated RNA molecules, and rely on the identification of the poly-(A) homopolymeric region to segment the barcode signal.

Recent works have shown that it is possible to sequence very short RNA reads (e.g. tRNAs, (Thomas et al. 2021; Lucas et al. 2023)), opening novel avenues to explore the small RNA (epi)transcriptome at single molecule resolution. To efficiently capture these short RNA molecules, adapted library preparation protocols, such as those used in ‘Nano-tRNAseq’, are required (Lucas et al. 2023). Notably, Nano-tRNAseq libraries substantially differ from standard DRS protocols, and the resulting reads contain very short poly-(A)-tails, which are RNA-DNA hybrids, and thus potentially lead to incorrect demultiplexing due to imprecise segmentation. Indeed, our initial explorations revealed that DeePlexiCon was not able to accurately demultiplex Nano-tRNAseq libraries, reaching 70% precision and 77% recall on Nano-tRNAseq libraries (**Table S6**).

To overcome this limitation, we specifically trained a new model to multiplex Nano-tRNAseq datasets (see **Fig. 3A**). Our results showed that SeqTagger was able to demultiplex up to four barcodes reaching high precision (98.3%) on the validation set (**Fig. 3B**), making it possible to multiplex Nano-tRNAseq libraries in the same flowcell, thus reducing Nano-tRNAseq library preparation and sequencing costs.

**Figure 3.**
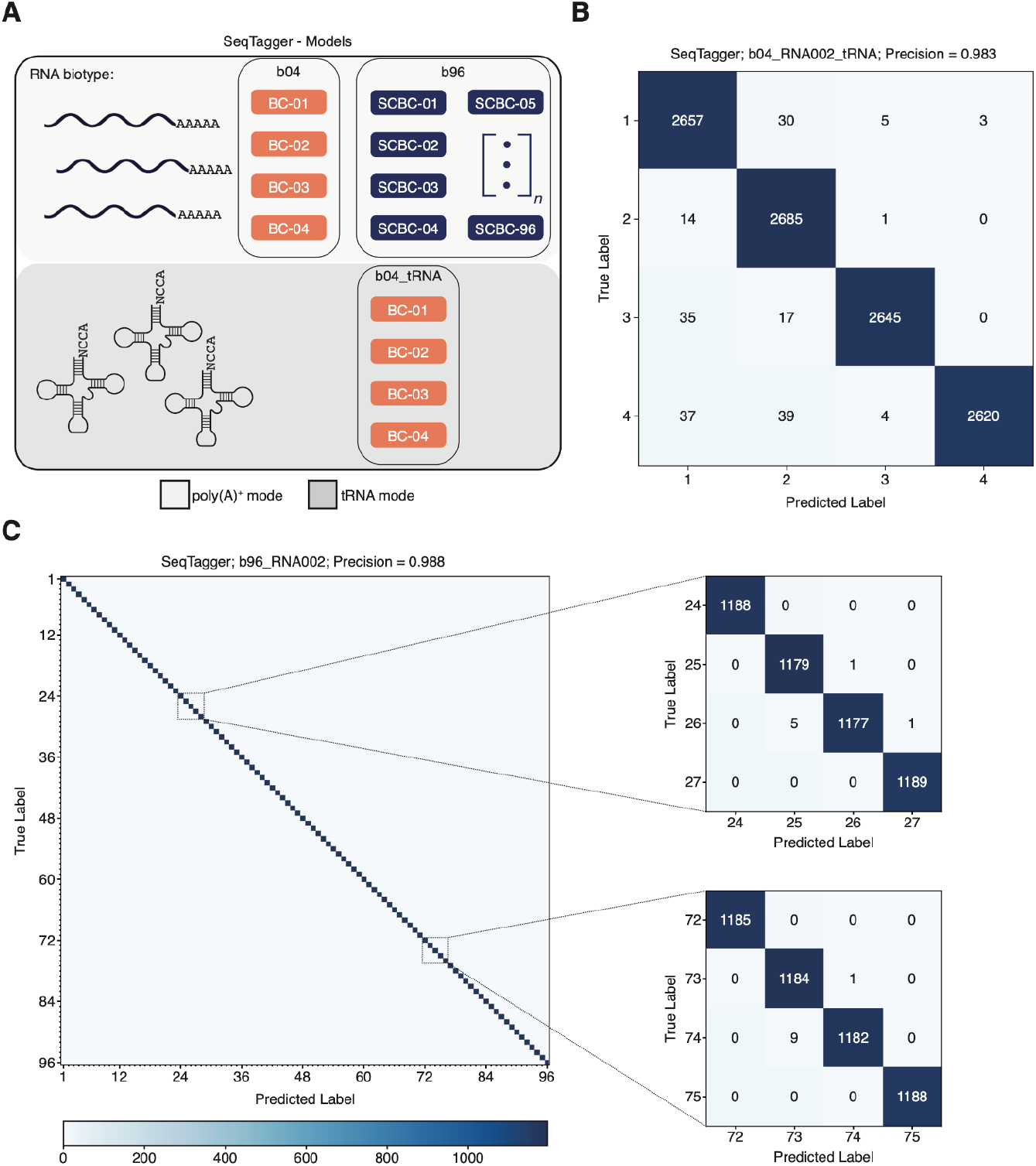
SeqTagger can be expanded to work on RNA-DNA hybrids (Nano-tRNAseq libraries) and with larger sets of barcodes. **(A)** Schematic overview of the demultiplexing models supported by SeqTagger. A four barcode (b04) and 96 barcode (b96) model are available. Additionally, a four barcode demuxing model is available for custom Nano-tRNAseq libraries (b04_tRNA). **(B)** Confusion matrix for the 4 barcode tRNA model (b04_tRNA) generated on Nano-tRNAseq validation data. Recorded precision is indicated on top. **(C)** Confusion matrix of the 96 barcode mRNA model (b96_RNA002) generated on the validation data. Recorded precision is indicated on top. Zoomed panels of the 96×96 confusion matrix for eight barcodes are shown on the right.

### SeqTagger can be extended to accurately demultiplex large barcode sets

Next, we wondered whether SeqTagger could be extended in terms of multiplexing capacity, i.e., by training a model that would predict additional barcodes, and whether demultiplexing performance would be significantly affected by the increased number of barcodes. To examine this, we trained a model containing a total 96 different barcodes (b96_RNA002), using the same procedure as previously described for four barcodes (**Fig. 3A**, *see also* **Supplemental Table S1**). Our results obtained on the validation dataset demonstrated that increasing the barcoding capacity by 24-fold only had a minor effect on model precision (98.8%) enabling simultaneous sequencing of up to 96 samples on a single flowcell (**Fig. 3C**).

We then examined the ability of SeqTagger to demultiplex *in vivo* samples using three independent test datasets. To this end, we extracted total RNA from human samples, which we poly(A)-tailed to make them amenable for direct RNA sequencing (*see* Methods). Next, we prepared three independent libraries barcoding each with one of the barcodes used for training the 96 barcode model (b96_RNA002). As expected, analysis of all three libraries revealed that the average base quality (baseQ) for barcodes present in the library was significantly higher than those absent, suggesting that base quality is an efficient parameter to remove false positive predictions (**Fig. 4A**, see also **Supplemental Fig. S2A**). Moreover, applying the default base quality filter used internally by SeqTagger (baseQ > 50), removed the majority of incorrectly assigned barcodes (96-97%) while maintaining between 93-95% of reads assigned to barcodes present in each library (**Fig. 4B**, see also **Supplemental Fig. S2B**). Of note, SCBC-25 which showed lower base quality on the independent test data (**Fig. 4A**, left panel), performed well on an additional independent test run containing this same barcode suggesting that the low performance on this particular test dataset was possibly related to the sample/library preparation, rather than the demultiplexing model (**Supplemental Fig. S2C**).

**Figure 4.**
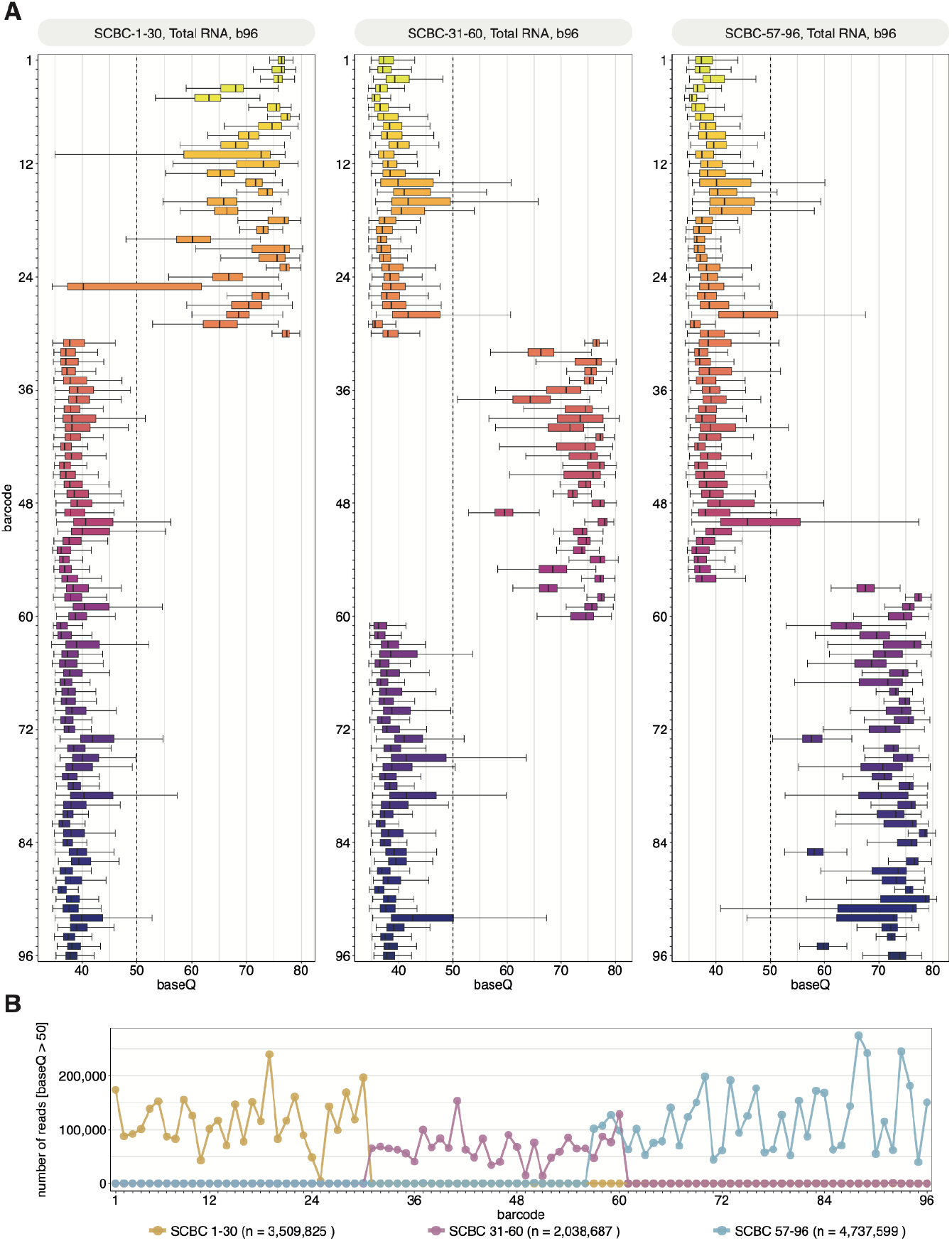
Performance of SeqTagger’s 96 barcode model on independent test data. **(A)** Boxplots depicting the base quality (baseQ) per barcode for three independent test runs using SCBC1-30 (left panel), SCBC-31-60 (middle panel) and SCBC-57-96 (right panel). Libraries were prepared using poly-(A)-tailed total RNA from human samples (*see* Methods). Boxes are limited by the lower quartile Q1 (bottom) and upper quartile Q3 (top). Whiskers are defined as 1.5 * IQR with outliers not shown. **(B)** Lineplot representing the total number of reads for each barcode (baseQ >50) for runs shown in Figure 4A, following demultiplexing. The total number of reads for each run with baseQ > 50 is indicated by *n*.

## DISCUSSION

Direct RNA nanopore sequencing has recently emerged as a transformative platform to characterise the (epi)transcriptome, offering unparalleled advantages in the analysis of RNA molecules, such as the possibility to detect its native RNA modifications (Garalde et al. 2018; Lucas and Novoa 2023). DRS has so far been used to address a wide range of biological questions in different biological systems, such as deciphering the order of intron removal during RNA splicing (Drexler et al. 2020), dissecting RNA modification dynamics upon chemical or biological stress (Huang et al. 2021; Begik et al. 2021; Delgado-Tejedor et al. 2023; Lucas et al. 2023), and elucidating viral RNA dynamics (Viehweger et al. 2019; Price et al. 2020; Kim et al. 2020; Baquero-Pérez et al. 2024), among others.

To date, commercial options for DRS multiplexing are lacking. Consequently, the field has relied on community-based solutions for demultiplexing, such as PorePlex (https://github.com/hyeshik/poreplex) or DeePlexiCon (Smith et al. 2020). The latter, while being accurate and widely adopted by the community (Rajan et al. 2023; Begik et al. 2021; Javaran et al. 2023; Gupta et al. 2023; White et al. 2023), suffers from several limitations: i) it currently supports only 4 barcodes; ii) to date, it only supports old sequencing chemistries (RNA002, currently being deprecated); iii) it requires CUDA v10 - and consequently won’t work with GPUs released after 2019, which use CUDA v11; iv) it does not work with custom reads that do not contain standard polyadenylated reads, such as Nano-tRNAseq reads, and v) the signal transformation step, is computationally expensive.

To overcome these limitations, we developed a novel approach for demultiplexing DRS reads, which relies on DNA basecalling of the barcode region, referred to as *SeqTagger*. We find that SeqTagger is a rapid and accurate demultiplexing program that is robust across different sequencing chemistries (RNA002, RNA004), devices (MinION, PromethION), and RNA species (mRNA, tRNA), supporting both fast5 and pod5 file formats. It achieves very high precision and recall on both smaller (4) and larger (96) barcode sets, thus opening exciting possibilities for the cost-effective sequencing of multiple samples on a single flowcell. To put these results into a broader context, a study published on cDNA demultiplexing on the ONT platform reached similar performance metrics on a twelve-barcode set compared to SeqTagger (b96_RNA002) (Wick et al. 2018). This suggests that demultiplexing using SeqTagger for direct RNA-sequencing can reach similar (RNA002) or higher (RNA004) classification metrics than current tools for cDNA demultiplexing.

We would like to note that the speed of demultiplexing with SeqTagger is still mostly bound by data access (**Supplemental Fig. S1A**) with our results reflecting a network filesystem. We observed that one million reads can be demultiplexed in 5 or 12 minutes when using a local SSD or HDD, respectively. Moreover, when data access isn’t a bottleneck, for example when the read signals are already loaded in system memory, SeqTagger can classify 50 barcodes in 5 milliseconds. This opens exciting possibilities such as real-time barcode classification of direct RNA sequencing reads, similar to what is currently available for DNA nanopore sequencing.

Finally, as an alternative approach to SeqTagger, one could imagine ligating either an RNA:DNA heteroduplex or an RNA:RNA barcode at the first step of library preparation (instead of DNA:DNA), which could be readily basecalled with existing RNA basecalling models and subsequently demultiplexed. However, there are several major challenges from both a wet- and dry-lab perspective with such an approach. Firstly, this would require a change to the library preparation protocol as ligases exhibit different substrate specificities, which in turn could reduce the overall efficiency of the library preparation (Bullard and Bowater 2006). Additionally, RNA oligonucleotides pose a substantial economic burden. For example, ordering only one strand of the oligonucleotides used in this study as an RNA molecule increases the cost of one RT adapter by ten fold, making large barcode sets prohibitively expensive. Moreover, RNA molecules are inherently less stable than DNA, which could lead to problems particularly when freeze-thawed multiple times, which is common for RT adapters. From a computational perspective, RNA oligonucleotides would require major changes to both the acquisition software MinKNOW and the basecalling software, as many definitions rely on differences in the current intensity signals obtained from RNA ligated to the DNA adapter.

## MATERIALS AND METHODS

### Plasmid extraction and linearization

We selected plasmids encoding *in vitro* transcripts from a T7 promoter that share little sequence similarity (*see* **Supplemental Table S7**) to enable unambiguous mapping of sequencing reads. Overnight cultures (LB-Ampicillin, 100µg/ml) containing *E*.*coli* transfected with the plasmids were processed using the Monarch® Plasmid DNA Miniprep Kit Protocol (NEB #T1010). Subsequently, plasmids were linearized using 25µl plasmid solution, 5µl of Enzyme 1, 5µl of Enzyme 2 (specific enzymes are mentioned in **Supplemental Table S7**), and 10µl of CutSmartBuffer in a total volume of 100µl for 3h at 37C. Volumes were topped up to 300µl using nuclease-free H_2_O before cleanup using 1x volume of basic Phenol:Chloroform:Isoamyl Alcohol (Sigma, P3803). Samples were vortexed followed by centrifugation at 16.000rpm for 5min. The aqueous phase was transferred to a new tube and supplemented with 0.1X 3M NaOAc (pH = 5.2), 2.5x EtOh abs., and 2µl of GlycoBlue™ (Thermo Fisher, AM9515). After overnight storage at −20C, precipitated plasmids were collected by centrifugation at 4C for 30min, 16000rpm. The pellet was washed twice with 70% EtOH and eluted in 30µl. Concentrations and purity were determined using a Nanodrop One/One^c^ (Thermo Fisher).

### In vitro transcription and poly(A)-tailing

*In vitro* transcription was performed using the AmpliScribe T7-Flash Transcription Kit (Biosearch Technologies, ASF3507) with 1µg of each linearized plasmid as input. The reaction was carried out as specified in the manufacturer’s instructions with an overnight incubation at 37ºC. The plasmid template was digested by adding 2µl of TURBO™ DNase (Thermo Fisher, AM2238) followed by incubation at 37C for 15min. *In vitro* transcripts were cleaned up using the RNeasy Mini Kit (Quiagen, 74104) and eluted in 60µl of nuclease-free H_2_O. Concentration and purity were determined using a Nanodrop One/One^c^ (Thermo Fisher). Transcript integrity was determined using a TapeStation 4150 (Agilent). For poly-(A)-tailing 2µl of 10X *E. coli* Poly(A) Polymerase Reaction Buffer, 2µl of 10mM ATP, 0.5µl SUPERase•In™ RNase Inhibitor (Thermo Fisher, AM2696) and 1µl of *E. coli* Poly(A) Polymerase (NEB, M0276S) were added to 7µg of input *in vitro* transcript in a total volume of 20µl. The reaction was carried out for 3min at 37C to obtain short poly-(A)-tails. Samples were cleaned up using the RNA Clean & Concentrator-5 Kit (Zymo Research, R1013) and eluted in 25µl of nuclease-free H_2_O.

### Barcode design

BC-01 to BC-04 correspond to barcodes previously described and used by DeePlexiCon (Smith et al. 2020). They are comprised of a 30nt long oligoA with a 5’P end, to enable ligation during library preparation and a 49nt long oligoB which contains a 10nt poly-d(T) 3’-end required for annealing to the target poly-(A)-containing library. SCBC-01 through SCBC-96 were designed to contain regions that are highly distinguishable in the sequence space (Doroschak et al. 2020). They are comprised of a 47nt long oligoA with a 5’P group to enable ligation during library preparation, and a 66nt long oligoB which contains a 10nt poly-d(T) 3’-end required for annealing to the target poly-(A)-containing library. See **Table S1** for details on all oligonucleotide sequences used to build barcoded libraries.

### Hybridization of custom RT adapters containing barcode sequences

All DNA oligos were ordered from IDT and annealed prior to library preparation using a final concentration of 1.4µM of each oligonucleotide, 0.01M Tris-HCl (pH = 7.5), and 0.05M NaCl in a total volume of 75µl nuclease-free H_2_O. The mixture was incubated at 94°C for 1 min and slowly cooled down (−0.1°C/sec) to room temperature. Small Aliquots were prepared to prevent repeated freeze/thawing of hybridized adapters. All oligonucleotides used in this work are listed in **SupplementalTable S1**.

### Direct RNA sequencing of training data

#### RNA002

Library preparation was carried out according to the manufacturer’s instructions (direct-rna-sequencing-sqk-rna002-DRS_9080_v2_revR_14Aug2019-minion) with some adjustments to enable barcode adapter ligation, prevent cross-contamination of barcodes and enable cost-efficient library preparation of multiple barcodes. Adapter ligation was carried out for 15min at room temperature using 100ng of input poly-A-tailed substrate, 0.5µl of custom barcode adapter (1.4µM), 1.5µl of NEBNext® Quick Ligation Reaction Buffer (NEB, B6058S), 0.5µl of RNAse Inhibitor (NEB, M0314S) and 0.75µl of concentrated T4 DNA Ligase (NEB, M0202T) in a total volume of 7.75µl nuclease-free H_2_O. Subsequently, 4µl of 5x SuperScript IV Buffer, 1µl of dNTPs (NEB, N0447S), 2µl 0.1 DTT, and 1µl of SuperScript™ IV Reverse Transcriptase (Thermo Fisher, 18090010) were added and the reaction incubated for 15min at 50C before heat inactivation for 2min at 70C. Reverse transcribed RNA:DNA hybrids were cleaned up using 1X RNAClean XP beads (Beckman Coulter, A63987), washed twice using freshly prepared 70% EtOH, and eluted in 3µl of nuclease-free H_2_O. 2.5µl of eluate was transferred to a new tube and RMX ligation was carried out individually for each barcode by adding 1µl of NEBNext® Quick Ligation Reaction Buffer (NEB, B6058S), 0.75µl of RMX adapter, and 0.375µl of T4 DNA Ligase (NEB, M0202T) in a total volume of 5µl nuclease-free H_2_O by incubating for 15min at room temperature. Libraries were cleaned up using 0.8X RNAClean XP beads (Beckman Coulter, A63987), and washed twice using 20µl of Wash Buffer (WSB) each time. Samples were eluted in 4µl of elution buffer (EB). Barcoded libraries were pooled at this step and the volume was adjusted to 18.75µl using nuclease-free H_2_O. Each library was mixed with 18.75µl of RNA running buffer (RRB) and run on a primed R9.4.1 flowcell using a MinION sequencer with MinKNOW acquisition software version v23.11.4. Live base calling and mapping were activated and runs stopped once ≥ 40.000 mapped reads were reached per barcode. Flowcells were reused multiple times following flushing (*see* Flowcell nuclease flushing).

#### RNA004

Library preparation was carried out according to the manufacturer’s instructions (direct-rna-sequencing-sqk-rna004-DRS_9195_v4_revB_20Sep2023-promethion). Changes to the above workflow for *RNA002* are mentioned, the remaining steps were identical. For adapter ligation 0.75µl of RNA Ligation Adapter (RLA) were used. Barcoded libraries were pooled and the volume was adjusted to 32µl using nuclease-free H_2_O. Before loading the libraries were mixed with 100µl of Sequencing Buffer (SB) and 68µl of Library Solution (LIS). Libraries were run on a primed FLO-PRO004RA flowcell using a PromethION 2 solo sequencer with MinKNOW acquisition software version v23.11.4.

### Flowcell nuclease flushing

Preventing carry-over during sequencing runs was paramount for generating high-quality training data. Hence, we adapted the existing flowcell flushing protocol (EXP-WSH004) by preparing a modified flowcell wash mix. To this end, we mixed 20µl of TURBO™ DNase (Thermo Fisher, AM2238) with 380µl of wash diluent (DIL, provided with EXP-WSH004). After loading the mix into the flowcell the reaction was incubated for 20 minutes. The remaining steps were identical to the manufacturer’s protocol. In our hands, this yielded negligible carry-over between runs (0.01-0.03%).

### Sample preparation and direct RNA sequencing of independent test data for b96

Total RNA was isolated using the AllPrep RNA/DNA/miRNA Universal Kit (Qiagen, 80224) using the manufacturer’s instructions. Frozen lung cryosections were homogenised with the TissueLyser mixer-mill disruptor (2 x 2 min, 25 Hz, Qiagen, Hilden, Germany). The quality of total RNA was assessed with an Agilent 2100 Bioanalyzer and Agilent RNA 6000 Nano Kit (Agilent Technologies, Boeblingen, Germany). Next, total RNA was split into a long (≥200nt) and short (<200nt) fraction using the RNeasy MinElute kit (Qiagen, #74204). Subsequently the long RNA fraction was poly-A-tailed (NEB, #M0276) in a total of 40µl. Individual poly-(A)-tailed samples were cleaned up using RNAClean XP beads (Beckman, #A63987).

For direct RNA-sequencing library preparation, individual samples (150 ng each) were ligated to pre-annealed (as described above) barcode-containing RT adapters (SCBCs) using concentrated T4 DNA Ligase (NEB, M0202T). Following reverse transcription using SuperScript™ IV Reverse Transcriptase (Thermo Fisher, 18090010), barcoded samples were pooled prior to clean-up RNAClean XP beads (Beckman, #A63987). For RMX ligation 250 ng of pooled library were used. Subsequent steps followed the standard direct RNA-sequencing library preparation protocol for RNA002 (direct-rna-sequencing-sqk-rna002-DRS_9080_v2_revS_14Aug2019-promethion). Libraries were run on a primed R9.4.1 flowcell using a PromethION sequencer with MinKNOW acquisition software version v23.11.4.

### Training the DNA basecaller

The training algorithm, *bonito (*https://github.com/nanoporetech/bonito/), was used to train the DNA basecaller model. Bonito requires as input signal chunks of fixed length and their corresponding barcode sequences. To know the signal-to-sequence correspondence for training purposes, one can either sequence one barcode per flowcell/run or ligate each barcode to a unique RNA molecule and later assign each read to the barcode by mapping reads to the corresponding references (see **Table S1** for barcode-sequence design).

We used 120,000 reads per barcode to train 4-barcode models (b04) and 40,000 reads per barcode for larger barcode sets (b96). Reads were randomly subsampled to reach the required number of reads per barcode. Importantly, we used only the last 3,000 (RNA002) or 2,000 (RNA004) samples of every barcode signal for training and basecalling. The reads with shorter barcodes were skipped.

The CTC-CRF models were trained with bonito v0.7.2. We used a window of 31 and a stride of 10. We tested three model versions with an increasing number of features used by the encoder (and model parameters): 96 (fast with 519,880 parameters), 384 (hac with 6,499,048 parameters) and 768 (sup with 24,793,192 parameters). Since we found only marginal differences in their accuracy for barcode basecalling (data not shown), we decided to use the lightest (and fastest) version with 96 features (fast).

### Basecalling of DNA barcode from direct RNA sequencing

The demultiplexing algorithm follows several steps: i) signal segmentation, ii) normalisation, iii) barcode sequence decoding, iv) barcode identification and v) quality filtering. First, the signal corresponding to the barcode is identified by T-test statistics in two window sizes (450 and 1,000) rolling over the first 30,000 read samples. We define barcode end as the position with the highest T-test score, additionally adding 100 samples of poly-A tail. Secondly, the barcode signal is normalised using median absolute deviation. Thirdly, the barcode sequence (and corresponding base qualities) are decoded using bonito v0.7.2 with a custom barcode basecalling model. Subsequently, the sequence is aligned onto barcode reference using minimap2 v2.26 executed with custom parameters: -k6 -w3 -A1 -B1 -O1 -E1 -c1 -m10 -s13. The best primary alignment for every barcode sequence is reported as the predicted barcode. Finally, only barcodes with median basecall quality above 50 are reported.

### Comparative analysis of demultiplexing software

For comparative performance analysis of demultiplexing software, we sampled 3 x 100,000 reads from an independent RNA002 run which consisted of IVT-01-04 ligated to BC-01-04 (**Fig. 2A**, *see also* **Supplemental Table S2**), which had not been used to train or validate the models. Each dataset was processed using the MasterOfPores (Cozzuto et al. 2020) version 3 nextflow (Di Tommaso et al. 2017) workflow, using either SeqTagger (model: *b04_RNA002*), DeePlexiCon (Smith et al. 2020) (model: *resnet20-final*.*h5*) or no demultiplexing (ground-truth) while keeping the remaining parameters identical. We compared default SeqTagger settings (baseQ ≥ 50, `-b 50`) with DeePlexiCon settings for high recovery (`-s 0.5`) and high accuracy (`-s 0.9`). Demultiplexing of individual samples (100,000 reads) was performed on a single GPU (RTX2080 Ti, CUDA10) and a single CPU with 12GB of memory allocated (`--granularity 25`). Samples were basecalled using guppy (v6.0.6, model = rna_r9.4.1_70bps_hac.cfg) and aligned to the reference sequence using minimap2 (v2.17) with `-ax map-ont -k14`.

Model performance metrics used to build confusion matrices and receiver operating characteristic curves were extracted using custom python scripts. Computational resources required per sample were extracted from the nextflow report (`-with-report`, see **Supplemental Table S4**) and analyzed using custom R scripts.

To compare the computation time of processes executed in the mop_preprocess workflow we used rep-1 of the benchmarking dataset and recorded the computation time required by each process (**Fig. 2C** bottom, and **Supplemental Fig. S1B)**. We realized that these results would underestimate the relative contribution of the mapping step as mapping was performed to a small reference of four sequences. To obtain a more realistic representation of the required computation time per process we sampled 100,000 reads from a poly-(A)-selected *M. musculus* library which was aligned to the mm39 reference genome (`minimap2, v2.17, -uf -ax splice -k14`). In addition, we activated counting, another optional step in the preprocessing pipeline. We obtained comparable results to the benchmarking dataset with DeePlexiCon taking up 40% of the overall computation time while SeqTagger reduces the time required to 8.5% (**Supplemental Fig. S1C**). To determine the performance of SeqTagger on *in vivo* datasets, we demultiplexed two runs containing poly(A)-tailed, total RNA extracted from *E*.*coli (Delgado-Tejedor et al*. *2023)* (**Supplemental Fig. S1D**, and **Supplemental Table S5**). Both runs were aligned to the *E. coli* rRNA reference (minimap2, v2.17, `-ax map-ont -k14`) after demultiplexing using either SeqTagger (model: *b04_RNA002*) or DeePlexiCon (model: *resnet20-final*.*h5*) with high recovery (-s 0.5) or high accuracy (-s 0.9) settings. Accuracy was determined by counting the number of reads assigned to each barcode in the fastq files (**Supplemental Table S5**). All scripts used to perform this analysis, as well as the reference sequences, can be found on the Github page (https://github.com/novoalab/SeqTagger/).

### Benchmarking model performance for RNA004, extended barcode models and tRNA models

To determine model performance on the new RNA004 chemistry we used an independent test dataset in which IVT-01-04 was ligated to BC-01-04, which hadn’t been previously used for model training or validation. Results were processed identically to the process described above (see *Comparative Analysis of demultiplexing software*) with exceptions mentioned next. Demultiplexing was run on a CUDA11-enabled single GPU (NVIDIA RTX A4000) and a single CPU with 12GB of memory allocated. For basecalling we used dorado (v0.5.3, https://github.com/nanoporetech/dorado, model = rna004_130bps_sup@v3.0.1). The extended barcode model (b96_RNA002) was trained on 24 runs each containing four *in vitro* transcripts ligated to the respective barcode. Confusion matrices were generated based on the validation dataset (analysis notebooks available at https://github.com/novoalab). The tRNA model was trained by using four individual Nano-tRNAseq runs performed on human cell lines, each using a different barcode sequence. Confusion matrices were generated based on the validation dataset. All runs used to generate training and testing data are listed in **SupplementalTable S2**.

## SOFTWARE AVAILABILITY

SeqTagger can be executed using a Docker container following the instructions on GitHub (https://github.com/novoalab/SeqTagger) or using the MasterOfPores (version 3.0) nextflow pipeline (MoP3) (https://github.com/biocorecrg/MOP3) (Cozzuto et al. 2023; Di Tommaso et al. 2017). Processing of raw fast5 files was performed using MasterOfPores (version 3.0). Full documentation on installation, usage, and dependencies related to MoP3 (including running SeqTagger via MoP3) can be found at https://biocorecrg.github.io/MoP3/.

## DATA ACCESS

Raw fast5 files and associated basecalled data in the form of bam files, generated in this work have been deposited in the European Nucleotide Archive (ENA) under accession code PRJEB78482. Basecalled fast5 datasets for poly-(A)-tailed, total RNA from *E. coli* used to determine cross-contamination are publicly available under accession code PRJEB42568. A summary of all datasets used in this work can be found in **Supplemental Table S2**.

## ACKNOWLEDGEMENTS

LPP was supported by funding from the European Union’s H2020 research and innovation program under Marie Sklodowska-Curie grant agreement No. 754422 and is currently supported by ERC funds (ERC-StG-2021 No 101042103 to EMN). GD is part of the ROPES ITN which received funding from the European Union’s Horizon 2020 research and innovation programme under the Marie Sklodowska-Curie grant agreement no. 956810. AD-T is supported by an FPI Severo-Ochoa fellowship by the Spanish Ministry of Economy, Industry and Competitiveness (MEIC). We would like to thank Dr. Huanle Liu for his insights and discussions at the commencement of this project. This work was supported by funds from the Spanish Ministry of Economy, Industry and Competitiveness (MEIC) (PID2021-128193NB-100 to EMN) and the European Research Council (ERC-StG-2021 No 101042103 to EMN). We acknowledge the support of the MEIC to the EMBL partnership, Centro de Excelencia Severo Ochoa, and CERCA Programme / Generalitat de Catalunya.

## AUTHOR CONTRIBUTIONS

GD generated training sets used to train SeqTagger models and performed bioinformatic analysis of the data, including comparative performance analysis to DeePlexiCon. LPP developed the SeqTagger algorithm and trained the models. LLL and RM contributed to generating nanopore sequencing datasets used to train the models. AD-T, LC and JP contributed to the data analysis and implementation of SeqTagger into MasterOfPores version 3. LPP and EMN conceived the project. EMN supervised the project. GD built the figures. GD, LPP and EMN wrote the manuscript, with contributions from all authors.

