## Supplemental_Figures for "SeqTagger, a rapid and accurate tool to demultiplex direct RNA nanopore sequencing datasets"

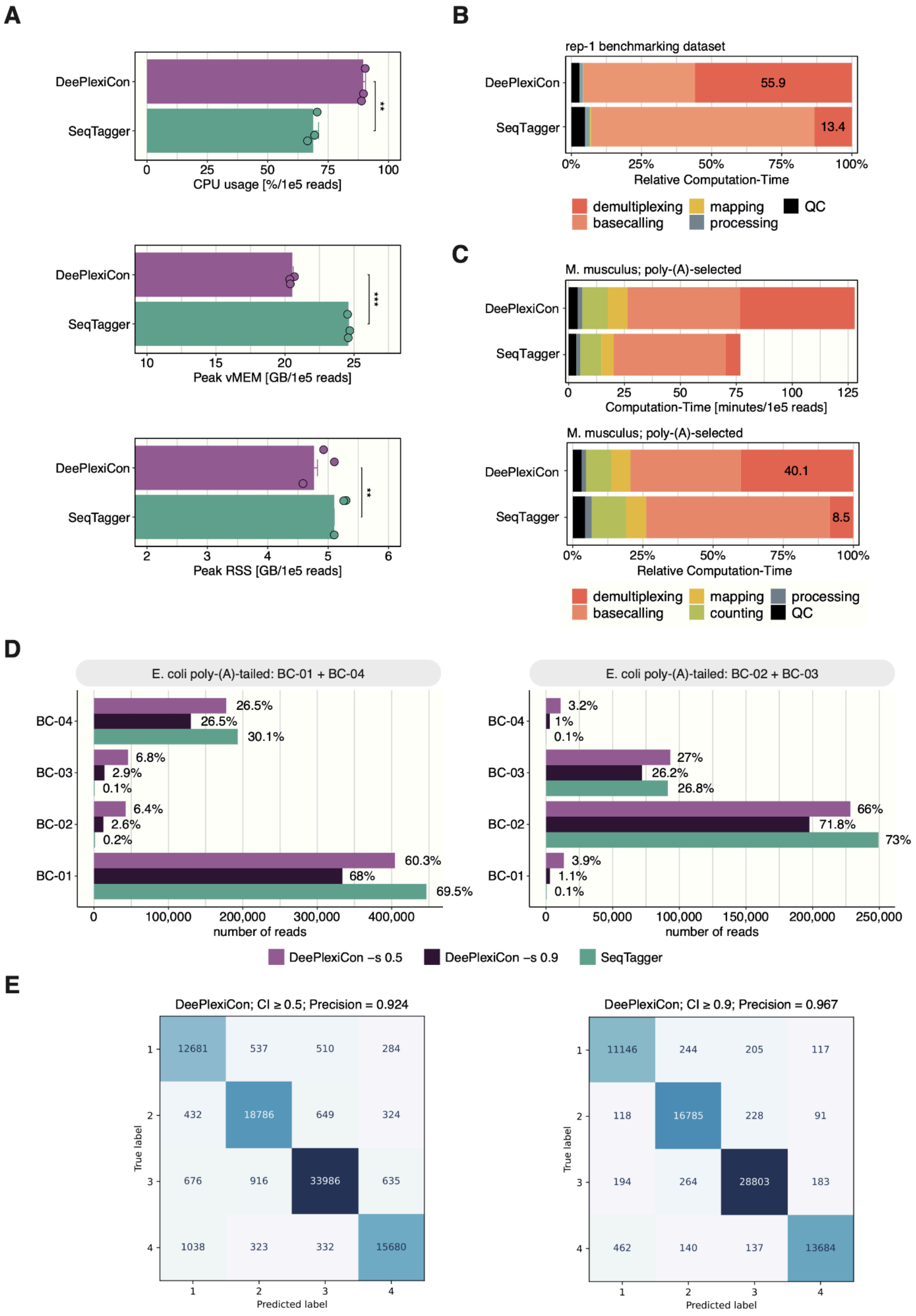
A
DeePlexiCon
SeqTagger
**
0
25
50
75
100
CPU usage [%/1e5 reads]
DeePlexiCon
SeqTagger
***
10
15
20
25
Peak vMEM [GB/1e5 reads]
DeePlexiCon
SeqTagger
**
2
3
4
5
6
Peak RSS [GB/1e5 reads]
B
rep-1 benchmarking dataset
DeePlexiCon
SeqTagger
55.9
13.4
0%
25%
50%
75%
100%
Relative Computation-Time
demultiplexing
basecalling
mapping
processing
QC
C
M. musculus; poly-(A)-selected
DeePlexiCon
SeqTagger
0
25
50
75
100
125
Computation-Time [minutes/1e5 reads]
M. musculus; poly-(A)-selected
DeePlexiCon
SeqTagger
40.1
8.5
0%
25%
50%
75%
100%
Relative Computation-Time
demultiplexing
basecalling
mapping
counting
processing
QC
D
E. coli poly-(A)-tailed: BC-01 + BC-04
BC-04
26.5%
26.5%
30.1%
BC-03
6.8%
2.9%
0.1%
BC-02
6.4%
2.6%
0.2%
BC-01
60.3%
68%
69.5%
0
100,000
200,000
300,000
400,000
number of reads
E. coli poly-(A)-tailed: BC-02 + BC-03
BC-04
3.2%
1%
0.1%
BC-03
27%
26.2%
26.8%
BC-02
66%
71.8%
73%
BC-01
3.9%
1.1%
0.1%
0
50,000
100,000
150,000
200,000
250,000
number of reads
DeePlexiCon –s 0.5
DeePlexiCon –s 0.9
SeqTagger
E
DeePlexiCon; CI ≥ 0.5; Precision = 0.924
True label
1
2
3
4
12681
537
510
284
432
18786
649
324
676
916
33986
635
1038
323
332
15680
1
2
3
4
Predicted label
DeePlexiCon; CI ≥ 0.9; Precision = 0.967
True label
1
2
3
4
11146
244
205
117
118
16785
228
91
194
264
28803
183
462
140
137
13684
1
2
3
4
Predicted label

**Supplemental Figure S1**. **Benchmarking results for computational requirements and model performance (A)** Barlpots depicting system requirements (CPU usage; peak vMEM; peak RSS) of DeePlexiCon and SeqTagger on the benchmarking dataset. Dots represent individual replicates with bars representing the mean value and error bars depicting +/-1 standard deviation. To determine statistical significance, a two-sided t-test was performed and results were corrected for multiple hypothesis testing using the Benjamini-Hochberg procedure. **(B)** Barplot depicting the relative contributions of each preprocessing workflow to the overall computation time on rep-1 of the benchmarking dataset. **(C)** Barplots depicting the absolute and relative contributions to the overall computation time of 100,000 reads sampled from a mouse poly-(A)-selected sample aligned to the mm39 genome. **(D)** Barplots representing the percentage of reads assigned to each barcode for two runs of total RNA from *E.coli* (poly-A-tailed). The first run (left) contained BC-01 and BC-04 while the second run (right) contained barcodes BC-02 and BC-03. Runs were demultiplexed with either SeqTagger (b04_RNA002) or DeePlexiCon (resnet20-final.h5) with high recovery (-s 0.5) or high accuracy (-s 0.9) settings (*see Methods*). **(E)** Confusion matrices corresponding to DeePlexiCon results for high recall (-s 0.5) and high precision (-s 0.9) on rep-1 of the benchmarking dataset.

---

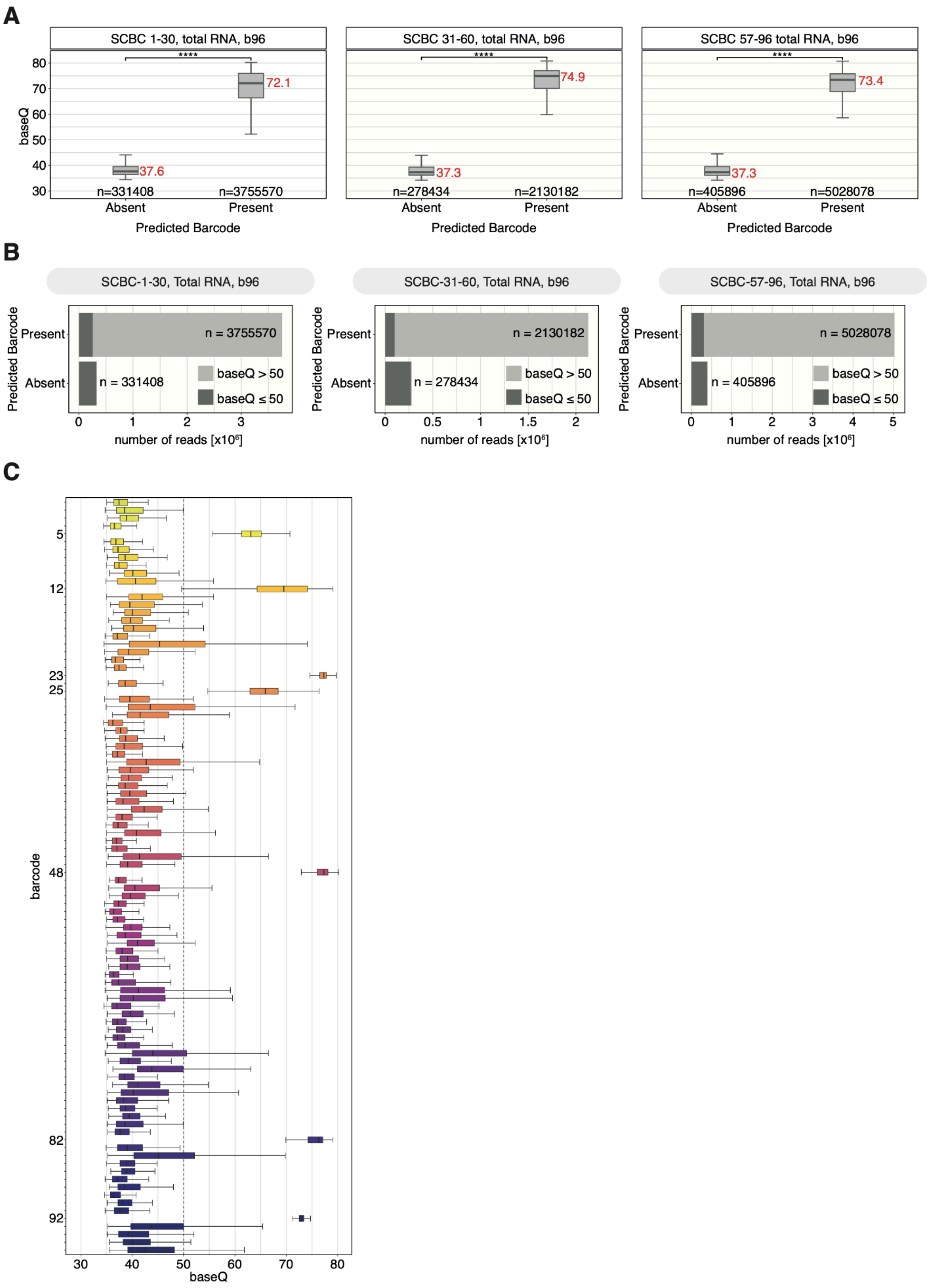

A
SCBC 1-30, total RNA, b96
SCBC 31-60, total RNA, b96
SCBC 57-96 total RNA, b96
baseQ
****
72.1
37.6
74.9
37.3
73.4
37.3
n=331408
n=3755570
n=278434
n=2130182
n=405896
n=5028078
Absent
Present
Predicted Barcode
B
SCBC-1-30, Total RNA, b96
SCBC-31-60, Total RNA, b96
SCBC-57-96, Total RNA, b96
Predicted Barcode
Present
Absent
n = 3755570
n = 331408
n = 2130182
n = 278434
n = 5028078
n = 405896
baseQ > 50
baseQ ≤ 50
number of reads [x10^6]
C
barcode
5
12
23
25
48
82
92
baseQ

**Supplemental Figure S2**. **Performance of SeqTagger's 96 barcode model on independent test data. (A)** Boxplots showing the baseQ distribution for barcodes present and absent in the three independent test runs. The Number of reads is indicated by *n* with the median value shown in red. Statistical analysis was performed using a two-sided non-parametric Wilcoxon test. Results were corrected for multiple-hypothesis testing using the Bonferroni procedure to obtain adjusted p-values (ns: p > 0.05, *: p <= 0.05, **: p <= 0.01, ***: p <= 0.001, ****: p <= 0.0001). **(B)** Barplots representing the total number of reads (*n*) for three independent test runs demultiplexed with SeqTagger's 96 barcode model (b96_RNA002). Colors indicate different baseQ thresholds. **(C)** Boxplots of base quality (baseQ) per barcode for an additional independent test run containing SCBC-05, SCBC-12, SCBC-23, SCBC-25, SCBC-48, SCBC-82 and SCBC-96. For Figures S2A and S2C the box is limited by the lower quartile Q1 (bottom) and upper quartile Q3 (top). Whiskers are defined as 1.5 * IQR with outliers not shown for visualization purposes.
